# Neurophysiological Dynamics of Metacontrol States: EEG Insights into Conflict Regulation

**DOI:** 10.1101/2024.09.28.615633

**Authors:** Xi Wang, Nasibeh Talebi, Xianzhen Zhou, Bernhard Hommel, Christian Beste

**Author notes:** <u>Address for correspondence</u> Bernhard Hommel, School of Psychology, Shandong Normal University, No. 88 East Wenhua Road, Jinan, 250014, Shandong Province, China. shared senior authorship (Bernhard Hommel, Christian Beste).

## Abstract

Understanding the neural mechanisms underlying metacontrol and conflict regulation is crucial for insights into cognitive flexibility and persistence. This study employed electroencephalography (EEG), EEG-beamforming and directed connectivity analyses to explore how varying metacontrol states influence conflict regulation at a neurophysiological level. Metacontrol states were manipulated by altering the frequency of congruent and incongruent trials across experimental blocks in a modified flanker task, and both behavioral and electrophysiological measures were analyzed. Behavioral data confirmed the experimental manipulation’s efficacy, showing an increase in persistence bias and a reduction in flexibility bias during increased conflict regulation. Electrophysiologically, theta band activity paralleled the behavioral data, suggesting that theta oscillations reflect the mismatch between expected metacontrol bias and actual task demands. Alpha and beta band dynamics differed across experimental blocks, though these changes did not directly mirror behavioral effects. Post-response alpha and beta activity were more pronounced in persistence-biased states, indicating a neural reset mechanism preparing for future cognitive demands. By using a novel artificial neural networks method, directed connectivity analyses revealed enhanced inter-regional communication during persistence states, suggesting stronger top-down control and sensorimotor integration. Overall, theta band activity was closely tied to metacontrol processes, while alpha and beta bands played a role in resetting the neural system for upcoming tasks. These findings provide a deeper understanding of the neural substrates involved in metacontrol and conflict monitoring, emphasizing the distinct roles of different frequency bands in these cognitive processes.

## Introduction

Action selection processes are central for daily life activities and have been investigated in cognitive neuroscience for decades. However, in the last years more research has been investigating the arbitration between different cognitive control strategies important during action selection. This arbitration, i.e. the decision which strategy is to employ during action selection to best cope with environmental demands, has been termed “metacontrol” (Hommel, 2015; Hommel and Colzato, 2017a; Mekern et al., 2019). According to the metacontrol state model (MSM) (Hommel and Colzato, 2017a), but also other framings of metacontrol (Beste et al., 2018; Goschke and Bolte, 2014), metacontrol varies between persistence and flexibility. A bias towards persistence reflects a focus on the current goal, strong competition between alternative stimulus or response representations, and the exclusive processing of task-relevant information, while a bias towards flexibility is characterized by a looser focus, weaker competition, and openness to nominally task-irrelevant information. There is increasing evidence that metacontrol biases vary systematically with the specific demands of the task at hand (Hommel and Colzato, 2017a; Mekern et al., 2019; Pi et al., 2024; Zhang et al., 2022).

One key function of human action control is the ability to deal with and resolve internal conflict. Conflict monitoring theory (Botvinick et al., 2001) posits a mechanism that monitors the occurrence of cognitive conflicts during information processing and serves to downregulate conflicts when needed. Cognitive conflict is often investigated in tasks that provide somewhat confusing information that is likely to activate mutually incompatible responses. For instance, in a flanker task, participants are presented with target symbols that are flanked by other, nominally task-irrelevant symbols that may signal the same response as the present target (a congruent trial) or another, competing response (an incongruent trial). Incongruent trials are assumed to create cognitive conflict that is detected and resolved by dedicated control mechanisms (Botvinick et al., 2001).

A recent study by Jia et al. (2024) was able to link conflict-induced control operations to metacontrol. It showed that incongruent trials induce a reduction of aperiodic EEG activity (Donoghue et al., 2020), also known as 1/f noise or scale-free activity (Buzsáki, 2006; He, 2014; Pertermann et al., 2019a, 2019b; Voytek and Knight, 2015), that extends into the next trial. Given that aperiodic activity has been suggested to reflect the current metacontrol state, with smaller degrees of activity reflecting a bias towards persistence and larger degrees reflecting a bias towards flexibility (Zhang et al., 2023), this suggests that cognitive/response conflict induces a metacontrol shift towards persistence. This fits with an observation of conflict-related behavioral adaptation (Gratton et al., 1992; Keye et al., 2013), showing that the effect of congruency on performance increases after congruent, and decreases after incongruent trials. This has been attributed to cognitive control operations that are triggered by the detection of internal conflict (Gratton et al., 1992; Keye et al., 2013), which would relax control after congruent trials but refresh control adjustments after incongruent trials. Control is stronger after incongruent trials, which in turn would reduce the congruency effect. In metacontrol terminology, relaxing and tightening cognitive control settings would correspond to shifts towards metacontrol flexibility and persistence, respectively. While the observation that aperiodic EEG activity is reflecting conflict-related changes of cognitive control increases our insights into the neurophysiological underpinnings of control operations (Van Schependom et al., 2024; Zhang et al., 2023), it raises the question how metacontrol is related to the much better investigated periodic activity. Numerous studies suggest important contributions of activity in theta, alpha and beta frequency bands to action control, even though the precise functional relevance of this activity is debated (Beste et al., 2023; Cavanagh and Frank, 2014; Cohen, 2014; Klimesch, 2012). Accordingly, the present study was guided by two research questions, namely, (i) whether periodic activity in different frequency bands is also modulated by metacontrol and (ii) which brain networks and directed communication therein are modulated by metacontrol states.

To induce a more persistence-biased versus a more flexibility-biased metacontrol state, we manipulated the frequency of congruent and incongruent trials in a flanker task. Frequent incongruent trials imply a rather frequent experience of response conflict, which should encourage participants to adopt a rather strict, persistent metacontrol state, to focus strongly on the goal and instruction, and to clearly distinguish between relevant and irrelevant information. Frequent congruent trials, in turn, imply a rather rare experience of conflict, which we thought should encourage a more lenient, flexible metacontrol state. We were interested to see whether these two metacontrol-relevant conditions would have differential effects on various frequency bands, and we were focusing in particular on theta, alpha, and beta band activity.

Theta band activity is possibly most important to metacontrol effects to become established. The reason is that (medial frontal) theta band activity is well-known to orchestrate different processes in distant brain regions (Buzsáki and Draguhn, 2004; Cavanagh and Frank, 2014) and is thus able to accomplish functions long-known to be essential for cognitive control (Miller and Cohen, 2001). Since metacontrol is very much about balancing whether new information becomes integrated or whether cognitive processes are shielded against this (Hommel, 2015; Hommel and Colzato, 2017a; Mekern et al., 2019), theta band activity likely is a natural candidate to also reflect metacontrol adjustments. Theta band activity has also been brought in connection with the dynamics important to integrate sensory and motor processes for the sake of goal-directed behavior (Beste et al., 2023). Interestingly, especially such integrated representations of perceptual and motor information have be suggested to be the central aspect upon which metacontrol dynamics unfolds (Hommel and Wiers, 2017). Moreover, theta band activity is also closely connected to broadband (gamma) activity (Roux and Uhlhaas, 2014; Uhlhaas, 2009) and it is this broadband activity contributes to the emergence to aperiodic activity (Gerster et al., 2022). All this leads to the hypothesis that especially theta band activity likely reflects metacontrol-related processes during conflict monitoring.

Yet, it needs to be noted that recent conceptualizations of theta band activity in the context of action control stress the importance of alpha and beta band activity (Beste et al., 2023). In this concept, theta band activity is thought to be affected by alpha and beta activity. Already this necessitates that also alpha and beta band activity is examined. Moreover, metacontrol is also about the adjustment of the processing of task-relevant and irrelevant information (Hommel and Colzato, 2017a; Mekern et al., 2019; Zhang et al., 2022). Such an adjustment may be implemented through inhibitory gating processes, known be reflected by alpha band activity (Klimesch, 2011; Klimesch et al., 2007), which also fits to conceptions of attentional selection processes reflected by this activity (Herrmann and Knight, 2001; Luck and Kappenman, 2013). However, according to the recent conception of the interplay of theta, alpha and beta band activity (Beste et al., 2023), it is again theta band activity that takes a central position in the interplay. Therefore, we hypothesize that all examined frequency bands show effects of the experimental manipulation, but that especially theta band activity takes a central role. If so, our two versions of the flanker task—FT_inc_ with frequent conflicts and a stronger persistence demand and FT_con_ with rare conflicts and more room for flexibility—should have an impact on and change activities in the frequency bands.

We adopted time-frequency and beamforming analyses to elucidate the roles of different frequency bands under our task manipulation, as well as the related source-level information. This is of relevance because current framings of the role of theta band activity during cognitive control is closely related to neuroanatomical considerations and an importance of medial frontal cortical regions (Cavanagh and Frank, 2014). The question to be explored is whether the regions that have been shown to play central roles in exerting cognitive control are also the regions involved in the dynamic adjustments of the cognitive control strategy (i.e. metacontrol). This is not necessarily the case, because the metacontrol can involve different strategies of how to adjust the cognitive control style (Hommel, 2015). Yet, it is likely that especially medial and inferior frontal regions, as well as superior parietal regions are involved. The reason is that these regions are known to exert a top-down control (i.e. frontal regions) (Aron et al., 2014; Miller and Cohen, 2001) and are involved in the integrated processing of sensor and motor aspects during action selection (i.e. parietal regions) (Beste et al., 2023; Gottlieb, 2007; Gottlieb and Snyder, 2010; Wilken et al., 2023; Yu et al., 2024). If two or more regions are involved for the different frequency bands examined, the question appears whether these brain regions exchange information (i.e. are connected). To answer this question, a nonlinear causal relationship estimation by artificial neural networks (nCREANN) was used (Elmers et al., 2024; Prochnow et al., 2024; Talebi et al., 2019).

## Methods

### Participants

*N*=43 native German speaking participants were tested in the current study. Five participants were excluded due to poor behavioral performance, EEG data quality and drop-out. The final sample consisted of *N*=38 participants (24 females; Mean age = 29.82, SD = 10.90 years). No psychiatric or neurological disorder was reported during the pre-study screening. All participants were provided written informed consent, and received either course credit or financial reimbursement in return for their participation. All participants were examined in three tasks (see below). The study was approved by the ethics committee of the Faculty of Medicine of the TU Dresden.

### Task Procedures

A standard flanker task, as previously used by our research group (Beste et al., 2010, 2007; Willemssen et al., 2009), was employed. In this task, we manipulated the frequency of congruent and incongruent trials to investigate our study question. A higher frequency of incongruent trials (70%; FT_inc_) was used to induce a persistence bias and a higher frequency of congruent trials (70%; FT_con_) to induce a flexibility bias. During the task, the participants sat in front of 20-inch TFT screen, on which the stimuli were presented. In each trial, a jittered fixation mark was presented first (700-1100ms). flanker stimuli were presented next for 200ms, followed by the target stimulus by 300ms (stimulus onset asynchrony =200 ms). Flanker and target stimuli disappeared simultaneously, and participants were required to respond within 650ms after onset of the target stimulus. During congruent trials, the flanker and the target stimuli pointed into the same direction; in incongruent trials, the flanker and the target stimuli pointed into the opposite directions, so as to induce a response conflict. Participants should press the left or right control key to indicate the direction of the target stimulus arrow. Depending on the block of the flanker task, the frequency of congruent and incongruent trials varied: In blocks with frequent incongruent trials (FT_inc_), the incongruent trials were presented in 70 percent and congruent trials 30 percent. In blocks with frequent congruent trials (FT_con_), the congruent trials were presented in 70 percent of all cases, and the incongruent trials 30 percent.

### EEG Data Acquisition and Pre-processing

EEG signals were recorded using QuickAmp and BrainAmp amplifiers (Brain Products GmbH, Gilching, Germany) with 60 Ag/AgCl electrodes embedded in an elastic cap (EasyCap Inc.). The software BrainVision Recorder 2.1 (Brain Products) was used for the recording. The sampling rate was 500 Hz, and the impedance of the electrodes was kept below 10 kΩ. The recorded data underwent preprocessing using the Automagic toolbox (Pedroni et al., 2019) and EEGLAB toolbox (Delorme and Makeig, 2004) on MATLAB R2022a (The MathWorks Corp.). The raw EEG signals were downsampled to 256 Hz and flat channels were removed before the data were re-referenced to an average reference. Next, we applied the PREP preprocessing pipeline (Bigdely-Shamlo et al., 2015) and the EEGLAB ‘clean_rawdata()’ pipeline. This sequence involved the removal of 50 Hz line noise and contamination by bad channels on the average reference. A finite impulse response high-pass filter (0.5 Hz, order 1286, stop-band attenuation -80 dB, transition band 0.25 - 0.75 Hz) was used to detect and remove flat-line, noisy, and outlier channels. Subsequently, to remove electromyographic artifacts, we employed a low-pass filter of 40 Hz (sinc finite impulse response filter; order: 86) (Widmann et al., 2015). A subtraction method was used for the removal of Electro-oculographic artifacts (Parra et al., 2005). A subsequent Independent Component Analysis was used to detect and exclude artifacts, such as muscle, cardiac, and remaining ocular artifacts based on Multiple Artifact Rejection Algorithm (MARA; Winkler et al., 2014, 2011). After pre-processing, continuous EEG signals were segmented into single trials for further time-frequency analysis. These EEG signals were locked to the target (e.g., where the middle triangle presenting) and segmented 2000ms before the target onset to 2000ms after the target onset. Besides, the time series of EEG signals in theta, alpha, and beta frequency bands were extracted using Hamming windowed sinc FIR filters. The filtered signals were then utilized for beamforming and connectivity analysis.

### Time-Frequency Analysis and Cluster-Based Permutation Testing

We utilized time-frequency (TF) decomposition through Morlet wavelets in the frequency domain for a between-conditions design. The wavelet length was set at three implicit Gaussian kernel standard deviations, with the number of cycles linearly ranging from three (for 3 Hz) to twelve (for 30 Hz). This process allowed for the estimation of average power in three relevant frequency bands (theta: 4–7 Hz, alpha: 8–12 Hz, and beta: 13–25 Hz) at each time point and EEG sensor. Baseline correction was performed using data from the time window spanning from -200ms to 0ms. Subsequently, we conducted a cluster-based permutation test using FieldTrip (Oostenveld et al., 2011) to analyze the time-frequency results of the three frequency bands of interest. This test aimed to compare differences between conditions: FT_inc_ vs. FT_con_ for both congruent and incongruent trials, as well as all incongruent trials vs. congruent trials. Utilizing the Monte Carlo approach, we generated 1,000 random draws to approximate the reference distribution of the permutation test. Clusters were considered significant if the corresponding p-values were below the critical alpha level of p = 0.025.

### Beamforming Analysis

To estimate directed functional connectivity from EEG signals, it is important to account for the volume conduction effect, which involves the transmission of electrical activity from the brain through conductive tissues (like the scalp and skull) and fluids (such as cerebrospinal fluid) to the electrodes placed on the scalp. Failure to address this effect can potentially distort the accuracy of connectivity estimates (Lai et al., 2018; Ruiz-Gómez et al., 2019). An approach commonly used to address this issue is to assess the connectivity of the underlying sources reconstructed from EEG recordings. In our study, we estimated the source time courses within each condition in the specified frequency bands using the linearly constrained minimum variance (LCMV) beamforming technique implemented in FieldTrip (Van Veen et al., 1997). Initially, a common spatial filter was derived from the time-locked average trials, which included concatenating all conditions. Subsequently, the time courses of the single trial source activities for each condition were computed using this common spatial filter. Clusters of theta, alpha, and beta activity within the LCMV-beamformed data were then identified by applying the Density-Based Spatial Clustering of Applications with Noise (DBSCAN) algorithm (Ester et al., 1996) to the Neural Activity Index (NAI) of the sources. In our investigation, we employed the DBSCAN method across different conditions, such as FT_inc_-inc, FT_inc__con, FT_con_-inc, and FT_con_-inc, for subsequent functional connectivity analysis. Additionally, we compared FT_inc_ with FT_con_ for both congruent and incongruent trials. Voxel selection was restricted to functional neuroanatomical regions exhibiting high activity, with the threshold set to the top 1% of the Neural Activity Index (NAI) distribution within labelled regions in the Automatic Anatomical Labeling (AAL) atlas (Tzourio-Mazoyer et al., 2002). The minimum cluster size was set to two voxels, with an epsilon value of 1.5 times the edge length of each voxel to identify neighbouring voxels. Selected clusters were visually inspected and chosen for further analysis based on voxel count and anatomical labels. The average of all voxel time courses within each selected cluster was considered as the activity estimate for that cluster in the connectivity analysis.

### Directed Functional Connectivity

To answer the aforementioned question regarding whether brain regions exchange information if they are involved for the different frequency bands, a connectivity analysis was carried out. No directed hypotheses about the connectivity pattern between sources of activity can be formulated, given the novelty of the research question focused in this study. Yet, directed communication between cortical modules is also key to cognitive control processes (Miller and Cohen, 2001), and it may similarly play a role in modulation by metacontrol. Therefore, and in exploratory data analyses, we examined the directional connectivity patterns in the theta, alpha, and beta frequency bands among the clusters identified by the DBSCAN algorithm through applying the nCREANN (nonlinear Causal Relationship Estimation by Artificial Neural Network) method (Talebi et al., 2019). The method employs an artificial neural network (ANN) to estimate directed connectivity among multiple regions using a nonlinear Multivariate Autoregressive (nMVAR) model. This model determines the current samples of brain regions by considering interactions among their past activities. Typically, the nMVAR model is utilized to illustrate temporal causality, where the past causes the future outcomes. The nCREANN method captures both linear and nonlinear dynamics of information flow among cortical areas, unlike standard linear approaches that solely focus on linear Multivariate Autoregressive (MVAR) models. Given that complex nonlinear behaviors exist in the nervous system from single neurons to the system level (He and Yang, 2021), linear methods risk oversimplifying the intricate dynamics of brain function. Indeed, nonlinear interactions play a crucial role in organizing information flow across cortical regions (Kodama and Galán, 2019; Yang et al., 2018). Evidence suggests the importance of both linear and nonlinear principles for a deeper understanding of neuro-dynamics at macroscopic levels (Chen et al., 2010; Cifre et al., 2021; Ferdousi et al., 2020; Friston, 2001; Nozari et al., 2020). In the nMVAR model, each signal’s current sample is represented as a (non)linear function of its previous values and the past values of other signals allowing for the inference of temporal causality. For an M-dimensional time series, a nonlinear MVAR model of order *p* is defined as

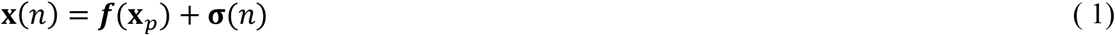

Where **X**_*p*_ = [*x*_*1*_(*n* – 1), *x*_2_(*n* – 1), …, *x*_*M*_(*n* – *p*)]^T^ is the vector of *p* past samples of (M) signals. The noise vector, ***σ***(*n*) = [σ_1_, σ_2_, …, σ_*M*_]^T^, is the model residual, and the quantitative explanation of how previous samples lead to the current values is provided by the nonlinear function ***f***(.). In the nCREANN method, the function ***f*** is split into linear and nonlinear parts:

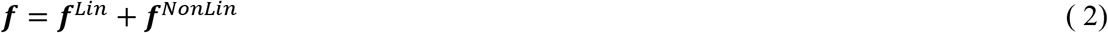

The Linear Connectivity (*lC*_*i*→*j*_) is determined using the ***f***^*Lin*^, to calculate the linear influence of *i*th region on the *j*th region, and the Nonlinear Connectivity, (*NC*_*i*→*j*_), is inferred based on the information embedded within ***f***^*NonLin*^, to establish the extent of the nonlinear effect of *x*_*i*_ on *x*_*j*_.

In our study, the nCREANN was applied to the time courses of the sources derived from LCMV in the FT_inc_-inc, FT_inc__con, FT_con_-inc, and FT_con_-inc conditions. We considered data points from trials occurring within the time interval of 0 to 1000ms after stimulus onset for connectivity analysis. To create a dataset of sufficient length for training the network, all trials were concatenated. Model order selection was determined using the Akaike and Schwartz criteria (Schneider and Neumaier, 2001), with a chosen order of *p* = 10) for all subjects and conditions. A single hidden layer feed-forward network with 10 hidden neurons, was trained in our study. The network took **X**_*p*_as input and aimed to predict **X**(*n*) as its output. The gradient descent error back-propagation (EBP) algorithm with momentum (α) and adaptive learning rate (η) was applied for the training. To enhance generalization, early stopping method was implemented. We employed a 10-fold permuted cross-validation technique, where the data was randomly split into 80% training, 10% validation, and 10% testing sets for each fold. Network parameters were updated in the ‘incremental’ mode, where each input presentation to the network prompted parameter updates. The input signals were scaled to be in the range [-1.5, 1.5], and the initial parameters were selected randomly within the range of [-0.5, 0.5].

We evaluated the nMVAR model’s fit using Mean Square Error (MSE) and the R-square values for both training and test data. R-Squared (R^2^), a key statistical measure in regression analysis, indicates the model’s goodness of fit, with values closer to 1 indicating better data fitting. MSE serves as a widely accepted metric for the assessment of the network performance. A well-trained network exhibits low training error, and the test error falls within the range of training error. Consistency between R^2^ values for training and test sets underscores the network’s robust generalization. To assess the significance of connectivity values, we employed a randomization test, generating 100 data sets through time-shifted surrogate techniques (Papana et al., 2013). This method preserves the dynamics of each time series while eliminating any causal effects between signals. The network parameters remained consistent with those used for the original data when applying nCREANN to the surrogate data. The connectivity patterns were visualized on a standard head model, with arrows indicating the flow of information from one cluster to another. Source cluster coordinates were obtained from the DBSCAN analysis (as described in the Beamforming Analysis section). Non-self-connections were plotted based on average values across all subjects. After obtaining all connections, we conducted paired comparisons for each connection between the metacontrol conditions to investigate which metacontrol condition exhibits higher connectivity. This analysis was performed using SPSS (Version 29.0) with Wilcoxon Signed-Rank Test. In addition, connectivity in reverse direction between each pair of clusters were also compared to determine the dominant direction.

## 3. Results

### 3.1 Behavioral results

A 2 × 2 repeated-measures ANOVA of Congruency and Frequency on the reaction time was performed (Figure 1). The results showed that the main effect of Frequency was not significant (*F*(1,37) = 1.44, *p* = .238, η_p_^2^ = .04), while the main effect of Congruency was significant (*F*(1,37) = 217.67, *p* < .001, η_p_^2^ = .86), with participants responding faster in congruent trials (339ms ± 7ms) than in incongruent trials (389ms ± 7ms). Most importantly, the Congruency × Frequency interaction was significant (*F*(1,37) = 246.19, p < .001, η_p_^2^ = .87), indicating that the congruency effect was substantially smaller in FT_inc_ blocks (22.46ms ± 3.19ms) than in FT_con_ blocks (78.59ms ± 4.44ms, *p* < .001) as shown in the Figure 1.

**Figure 1.**
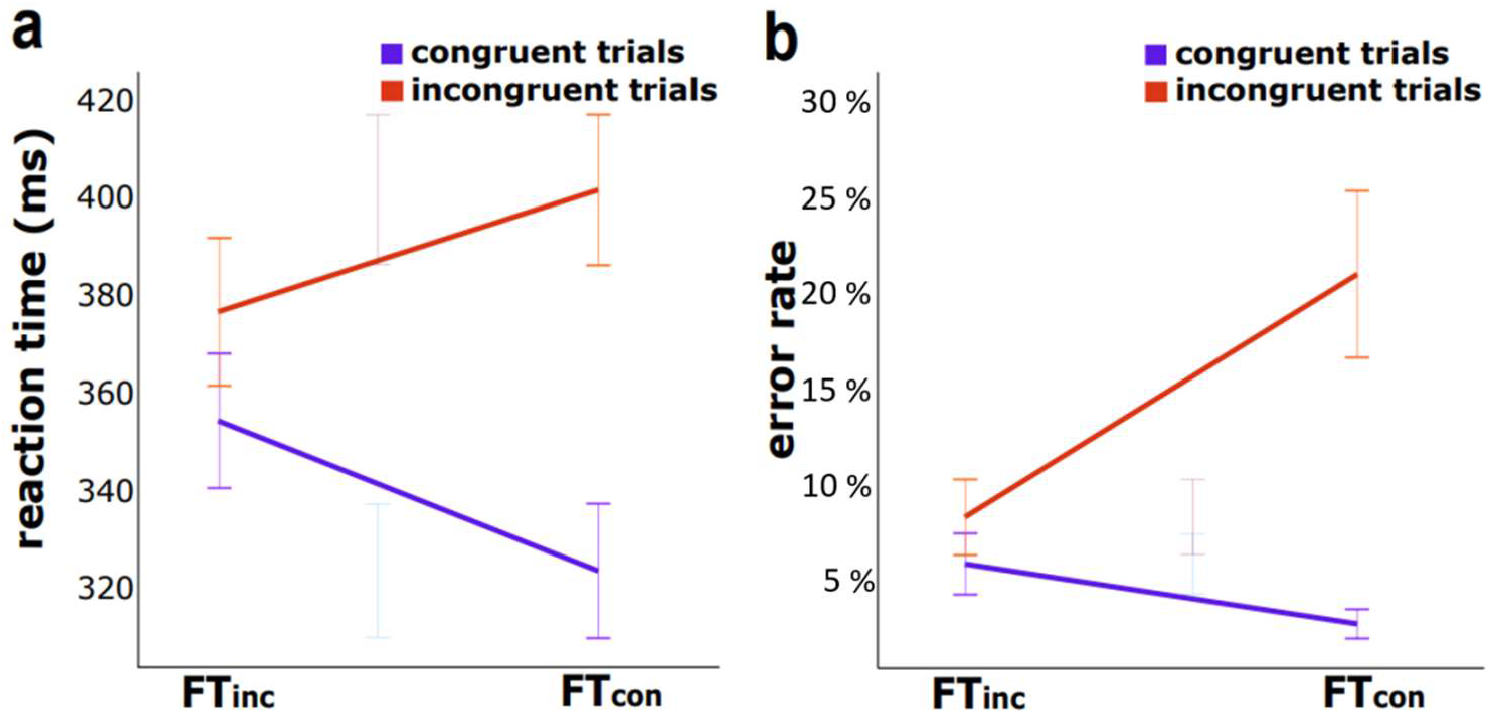
(a) Reaction time of FT_con_ and FT_inc_ conditions in congruent (purple line) and incongruent (orange line) trials. (b) Error rate of FT_con_ and FT_inc_ conditions in congruent and incongruent trials. Error bars represent standard deviations.

A 2 × 2 repeated-measures ANOVA of Congruency and Frequency on the error rates was performed (Figure 1). The results revealed significant main effect of Frequency (*F*(1,37) = 41.14, *p* < .001, η_p2_ = .53), with higher error rates in FT_con_ (11.8% ± 1.2%) than in FT_inc_ blocks (7% ± 0.8%). The main effect of Congruency was also significant (*F*(1,37) = 72.90, *p* < .001, η_p_^2^ = .66), with higher error rates in incongruent trials (14.6% ± 1.5%) than in congruent trials (4.2% ± 0.5%). Importantly, a significant Frequency × Congruency interaction (*F*(1,37) = 82.75, p < .001, η_p_^2^ = .69), showed that the congruency effect was substantially smaller in FT_inc_ blocks (2% ± .01%) than in FT_con_ blocks (18% ± 2%, *p* < .001) as shown in the Figure 1.

### 3.2 Time frequency and beamforming of metacontrol effects analysis

At the sensor level, the cluster-based permutation tests revealed significant clusters for metacontrol effects during different congruency trials on all three frequency bands. At the source level, the LCMV beamforming and DBSCAN were used to localize the generator of theta band activity (TBA), alpha band activity (ABA) and beta band activity (BBA) for what we found at the sensor level results.

#### 3.2.1 Congruent trials

For the theta band, there is a positive activity shows that FT_inc_ has a higher activity than FT_con_ (Table 1, Figure 2a). For the alpha band, a negative activity was found, for which FT_inc_ has a lower activity than FT_con_. A negative activity was also found for the beta band, for which FT_inc_ has a lower activity than FT_con_. The DBSCAN revealed that the positive cluster for the theta band was located at right inferior frontal extending to postcentral (Table 1, Figure 2a). For the alpha band, three negative clusters located at 1) right inferior frontal, 2) left angular, 3) right angular extend to superior parietal. For the beta band, three negative clusters located at 1) left middle occipital, 2) left precentral and postcentral, 3) right precentral and postcentral.

**Table 1.** Time Frequency results for the FT_inc_ - FT_con_ comparison respectively for congruent and incongruent trials. TBA, theta band activity. ABA, alpha band activity. BBA, beta band activity. Significant time windows are shown relative to the target display.

| Metacontrol effect congruent trials |  |  |  |  |
| --- | --- | --- | --- | --- |
| Frequency band | Comparison | P value | Significant time window | DBSCAN clusters |
| TBA | $FT_{inc} > FT_{con}$ | $p < .001$ | 141-926ms | right inferior frontal extending to postcentral (rIF-PC, BA44, BA1/2/3/4) |
| ABA | $FT_{inc} < FT_{con}$ | $p < .023$ | 375-715ms | right inferior frontal (rIF, BA45/47), left angular (lA, BA39), right angular extend to superior parietal (rA-SP, BA39, BA5) |
| BBA | $FT_{inc} < FT_{con}$ | $p < .006$ | 352-703ms | left middle occipital (lMO, BA18/19), left precentral and postcentral (lPPC, BA1/2/3/4), right precentral and postcentral (rPPC, BA1/2/3/4) |
| Metacontrol effect incongruent trials |  |  |  |  |
| TBA | $FT_{inc} < FT_{con}$ | $p < .001$ | 0-691ms | left inferior temporal (lIT, BA20), right middle temporal to superior temporal (rMT-ST, BA21/22) |
| ABA | $FT_{inc} < FT_{con}$ | $p < .005$ | 188-469ms | right middle temporal (rMT, BA10), left superior frontal to supplementary motor area (lSF-SMA, BA10/11/12, BA6) |
| BBA | $FT_{inc} < FT_{con}$ | $p < .001$ | 118-996ms | right hippocampus and parahippocampus (rH-PH, BA34/35), right inferior occipital (rIO, BA17/18/19) |

**Figure 2.**
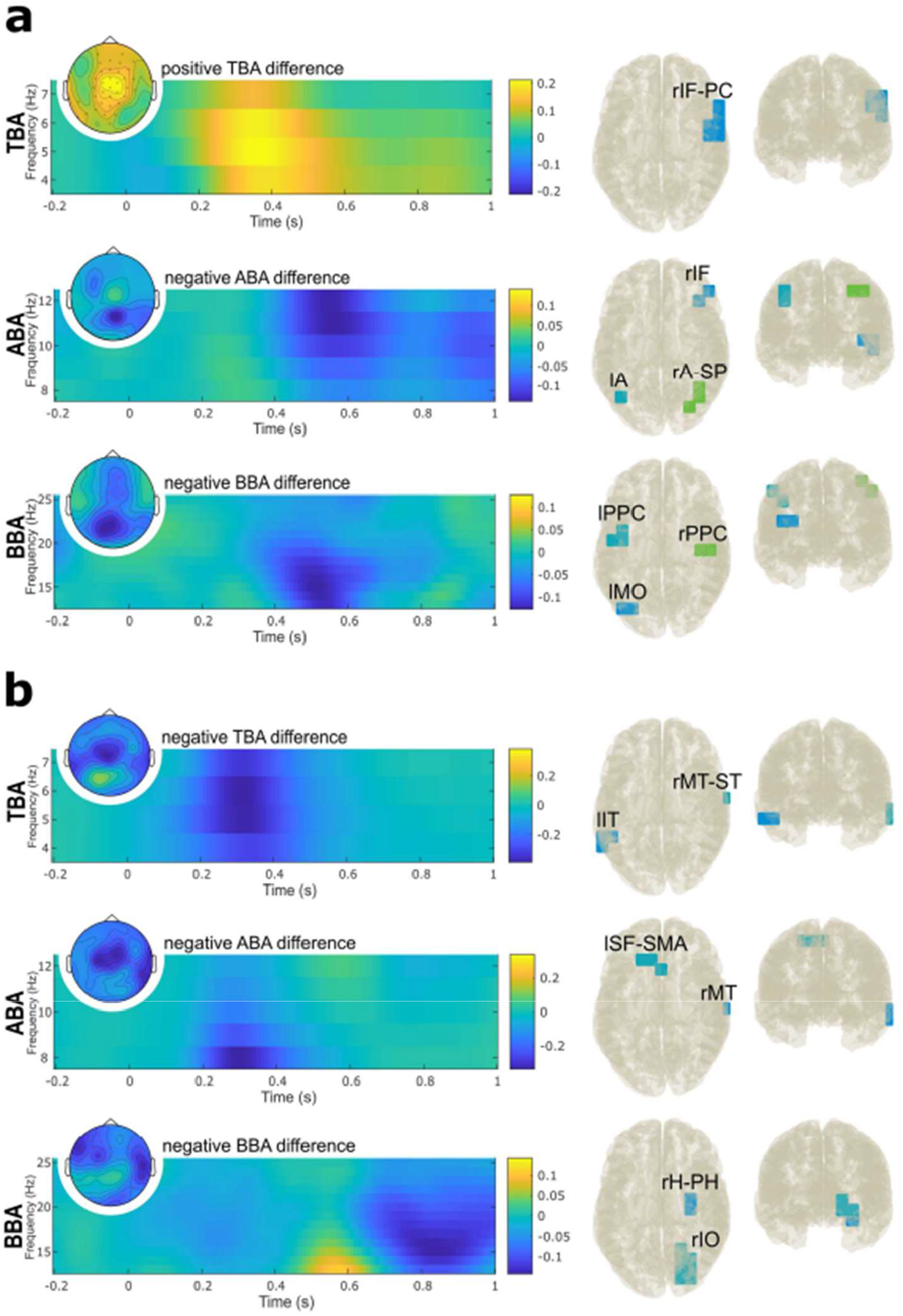
(a) Congruent Trials. Left panel, results of the time-frequency decomposition (TF plots) and the cluster-based permutation testing (topographic plots) for FT_inc_ and FT_con_ contrast in congruent trials. TF plots show the mean power difference across significant electrodes for each frequency band, as identified by cluster-based permutation testing. Topographic plots depict the average power difference within significant time windows for each frequency band, identified through cluster-based permutation testing. Right panel, results of DBSCAN for FT_inc_ and FT_con_ contrast in congruent trials. The first column displays top view and second column displays back view. Right upper panel, rIF-PC = right inferior frontal extending to postcentral. Right middle panel, rIF = right inferior frontal, lA = left angular, rA-SP = right angular extend to superior parietal. Right lower panel, lMO = left middle occipital, lPPC = left precentral and postcentral, rPPC = right precentral and postcentral. (b) Incongruent Trials. Left panel, results of the time-frequency decomposition and the cluster-based permutation testing for FT_inc_ and FT_con_ contrast in incongruent trials. Right panel, results of DBSCAN for FT_inc_ and FT_con_ contrast in incongruent trials. Right upper panel, lIT = left inferior temporal, rMT-ST = right middle temporal to superior temporal. Right middle panel, rMT = right middle temporal, lSF-SMA = left superior frontal to supplementary motor area. Right lower panel, rH-PH = right hippocampus and parahippocampus, rIO = right inferior occipital.

#### 3.2.2 Incongruent trials

Negative activity was found for all TBA, ABA and BBA. The same pattern revealed that FT_inc_ has a lower activity than FT_con_ (Table1, Figure 2b). The DBSCAN revealed a different pattern for incongruent trials (Table1, Figure 2b). For the theta band, two negative clusters located at 1) left inferior temporal, 2) right middle temporal to superior temporal. For the alpha band, two negative clusters located at 1) right middle temporal, 2) left superior frontal to supplementary motor area. For the beta band, two negative clusters located at 1) right hippocampus and parahippocampus, 2) right inferior occipital.

Based on the results of the DBSCAN, the directed connectivities were calculated using nCREANN. Since especially the effects of the FT_inc_ compared to FT_con_ experimental conditions are of interest in this study, these effects for congruent (section 3.3) and incongruent trials (section 3.4) are outlined below. Other results of the connectivity analysis are shown in the Supplemental material: Supplementary Figure S1 outlines the pattern of linear and nonlinear connectivities in the FT_inc_ compared to FT_con_ for congruent trials in the theta, alpha and beta frequency band. Statistics of the linear connectivity directions are provided in Supplemental Table S1, statistics of the nonlinear connectivity directions are provided in Supplemental Table S2. Supplementary Figure S2 outlines the pattern of linear and nonlinear connectivities in the FT_inc_ compared to FT_con_ for incongruent trials in the theta, alpha and beta frequency band. Statistics of the linear connectivity directions are provided in Supplemental Table S3, statistics of the nonlinear connectivity directions are provided in Supplemental Table S4.

### 3.3 Metacontrol effects on connectivity directions of congruent trials

As shown in Supplemental Figure S1, the three cluster of activity obtained for the alpha frequency band (see Figure 2a) revealed linear and nonlinear connections with each other. The same was the case for the beta frequency band.

Non-parametric tests were conducted for each connectivity direction to examine differences between FT_inc_ compared to FT_con_ experimental conditions – that is the metacontrol effect. The linear connectivity showed significant differences are shown as follows (Table 2, Figure 3): For the theta band, no linear connectivities could be calculated, because there was only a single cluster of activity. For the alpha band, the linear connectivity from lA to rIF, rA-SP to rIF, and rA-SP to lA were significantly higher in FT_inc_ compared to FT_con_. For the beta band, lMO to lPPC connectivity is higher in FT_inc_ compared to FT_con_. Other linear connectivity measures did not show significant differences. The nonlinear (Table 2, Figure 3), and for the alpha band, the connectivity from rA-SP to rIF was higher in FT_inc_ than in FT_con_. For the beta band, rPPC to lMO nonlinear connectivity is higher in FT_con_ compared to FT_inc_. Other nonlinear connectivity measures did not show significant differences.

**Table 2.**
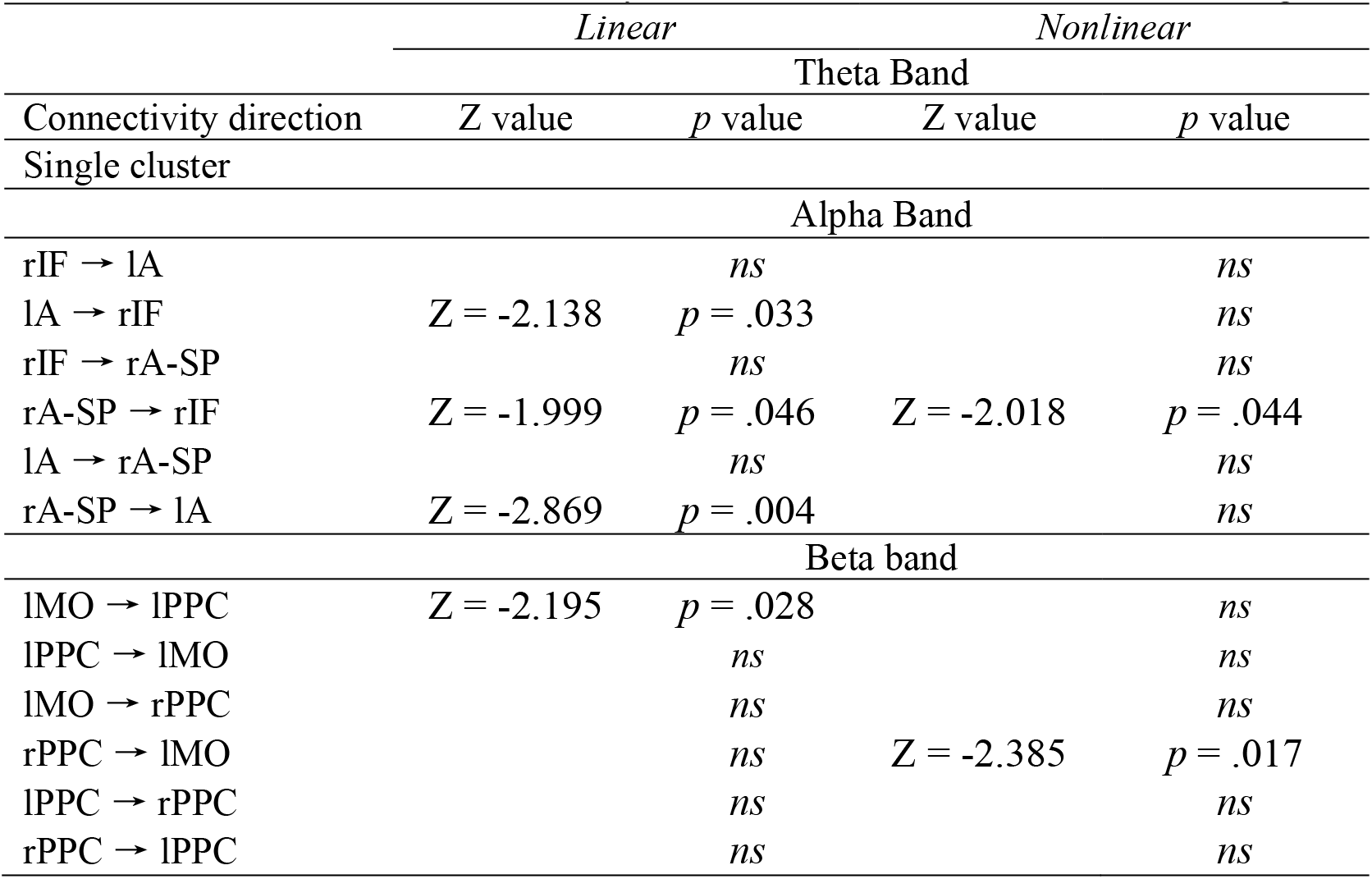
Linear and nonlinear connectivity directions between FT_inc_ and FT_con_ of congruent trials.

**Figure 3.**
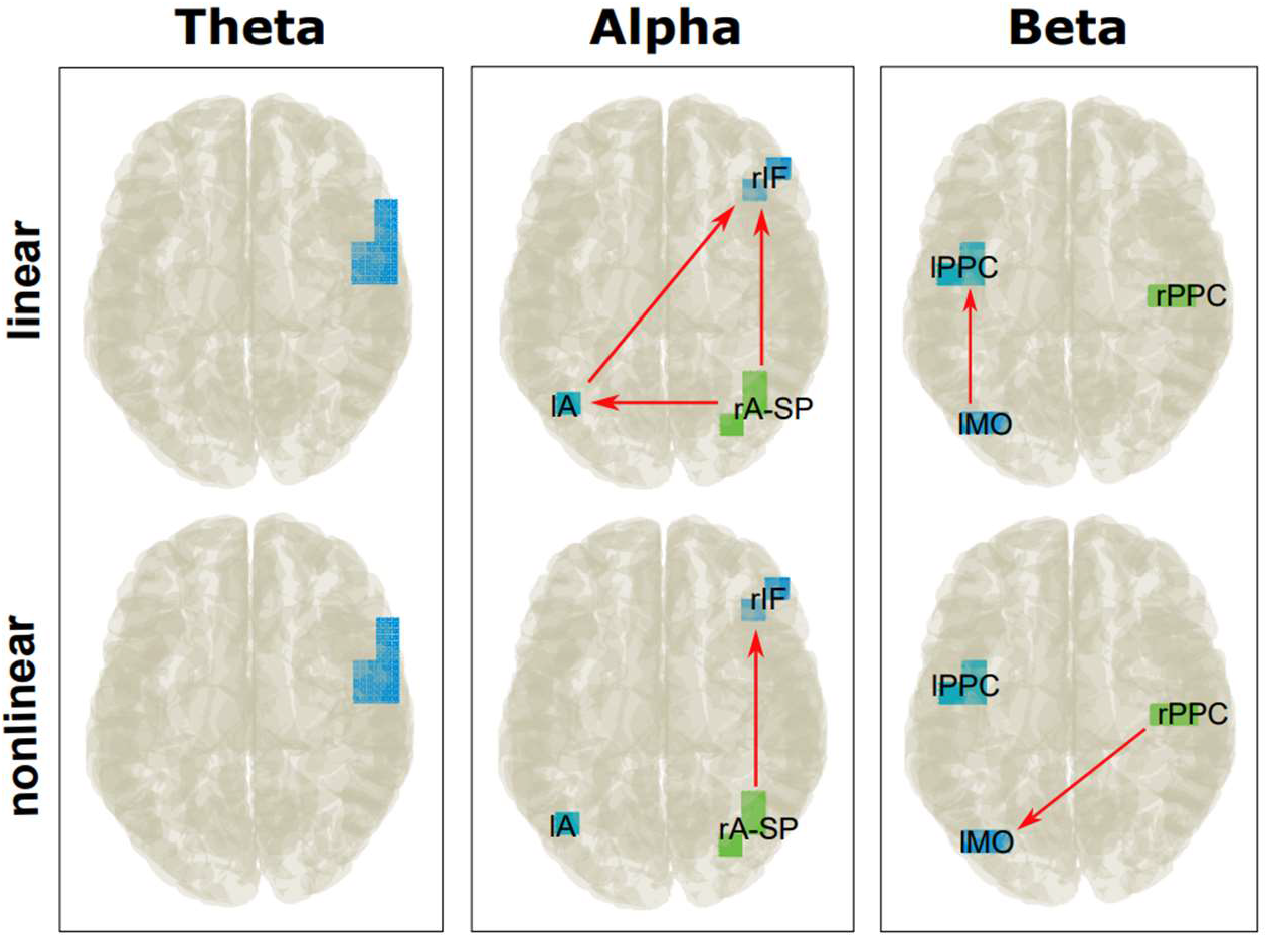
The nCREANN Linear and Nonlinear connectivity between FT_inc_ and FT_con_ of congruent trials after non-parametric tests. The presented view is top view. Red arrows indicate a significant difference in the connectivity between FT_inc_ and FT_con_.

### 3.4 Metacontrol effects on connectivity directions of incongruent trials

As shown in Supplemental Figure S2, the clusters of activity obtained for all frequency bands (see Figure 2b) revealed linear and nonlinear connections with each other.

Non-parametric tests were conducted for each connectivity direction to examine differences between FT_inc_ compared to FT_con_ experimental conditions – that is the metacontrol effect. The linear connectivity showed significant differences (Table 3, Figure 4): For the theta band, no linear connectivity measure showed significant difference. For the alpha band, lSF-SMA to rMT connectivity is higher in FT_con_ compared to FT_inc_., For beta band, no linear connectivity measure showed significant difference. Regarding the nonlinear connectivity (Table 3, Figure 4), the theta band revealed that the connectivity from the rMT-ST to the lIT is higher in FT_con_ than in FT_inc_. For the alpha and beta band, no nonlinear connectivity showed significant differences.

**Table 3.**
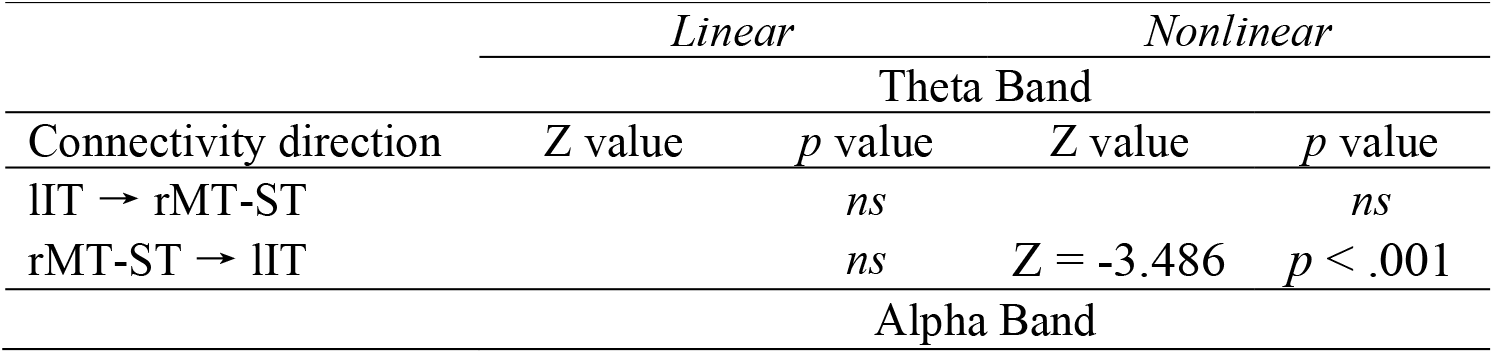
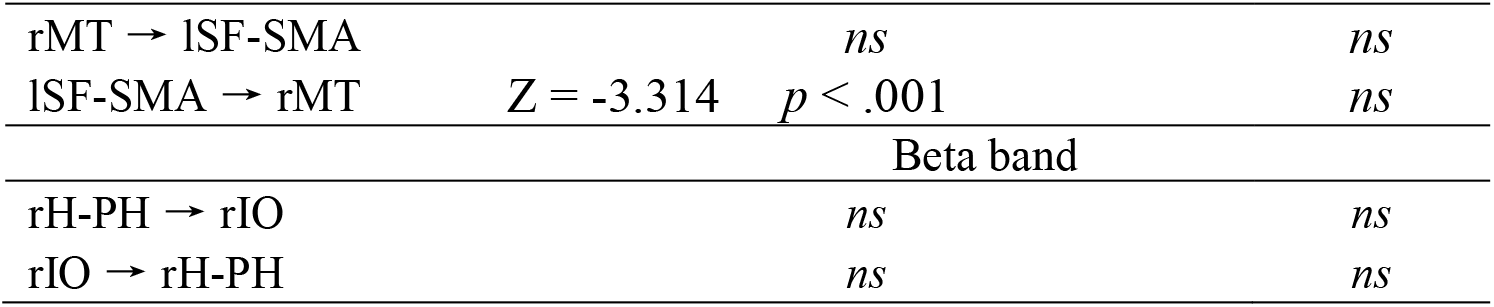
Linear and nonlinear connectivity directions between FT_inc_ and FT_con_ of incongruent trials.

**Figure 4.**
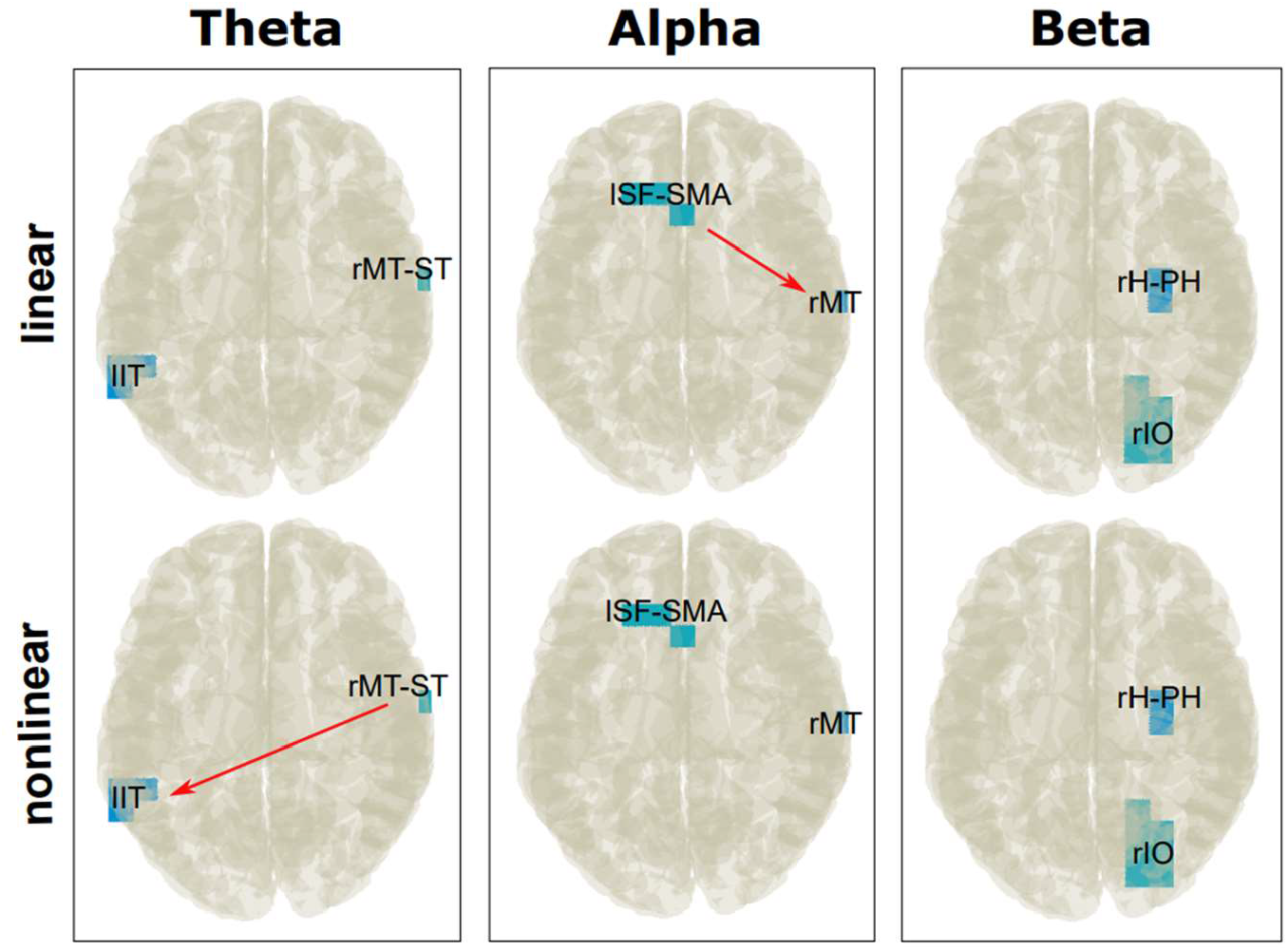
The nCREANN Linear and Nonlinear connectivity between FT_inc_ and FT_con_ of incongruent trials after non-parametric tests. The presented view is top view. Red arrows indicate a significant difference of the connectivity between FT_inc_ and FT_con_.

## Discussion

In the current study we examined the effects of metacontrol states on conflict regulation by means of EEG methods. To manipulate metacontrol states, we experimentally varied the relative frequency of congruent and incongruent trials in separate experimental blocks and tested the effect of this manipulation on the processing of congruent and incongruent trials. The study provides insights as to how metacontrol affects the processing of response conflicts and which electrophysiological processes including the directed communication between functional neuroanatomical structures are modulated.

The behavioral data show a clear, expected effect of the experimental manipulation: The congruency effect (i.e. the difference between congruent and incongruent trials) in response time and accuracy was larger if congruent trials were frequent (FT_con_) than if incongruent trials were frequent (FT_inc_). This pattern is common in studies where the relative frequency of congruent and incongruent trials is varied and has also been reported from other tasks that induce response conflict, like the Stroop task (Logan and Zbrodoff, 1979) or the Simon task (Hommel, 1994). From a metacontrol perspective, such patterns represent a stronger bias towards metacontrol persistence if the probability of (response) conflict is high and a stronger bias towards metacontrol flexibility if the probability of (response) conflict is low. Our findings are therefore in line with our expectation that the FT_inc_ block induced a more persistent state while the FT_con_ block induced a more flexible state. The data thus show that metacontrol can be examined through a manipulation of the frequencies of occasions requiring increased or moderate levels of cognitive control. The metacontrol bias towards persistence or flexibility can therefore be seen as a direct function of the likelihood to engage in control-demanding cognitive processes. Given these indications that our main manipulation worked as expected, we now can turn to the question how it affected periodic EEG activity.

### Modulations of theta band dynamics

The theta frequency band showed a pattern paralleling that observed for the behavioral data, and the impact of our metacontrol manipulation was clearly visible before the mean reaction time. More specifically, theta band activity increased more during congruent trials in the FT_inc_ block than during congruent trials in the FT_con_ block (Table 1, Figure 2). Incongruent trials showed the reversed pattern with stronger theta band activity increases in the FT_con_ block than in the FT_inc_ block (Table 1, Figure 2). Current views on the role of theta band activity suggest that it represents a response to a surprise signal and indicates the need to increase cognitive control (Beste et al., 2023; Cavanagh and Frank, 2014). Usually, cognitive control does not need to be increased in congruent trials. So, how can the pattern of findings be reconciled with current concepts of theta band activity?

The idea behind introducing blocks with different frequencies of congruent and incongruent trials was to establish a context biasing metacontrol towards persistence or flexibility. Within such a persistence-prone context, congruent trials are rare and therefore produce a surprise. The same is the case for incongruent trials presented in a flexibility-prone context. Therefore, it seems that it is the mismatch or conflict between what is to be expected based on the actual metacontrol bias and the processing required in the actual situation is reflected by the modulations of theta band activity. In other words, theta band activity is sensitive to metacontrol processes and the expectations they imply. However, interpretations of theta as signaling a need for control strongly relate to the medial frontal cortex (Cavanagh and Frank, 2014), but not the regions in the ventral processing stream of the visual system and the right inferior frontal cortex as found in the current study. According to metacontrol theories, a bias towards persistence or flexibility implies a stricter or a looser focus on the processing of task-relevant information (Hommel and Colzato, 2017a; Mekern et al., 2019), which can emerge through the modulation of attentional processes (Hommel and Colzato, 2017b). Although the rIFG has generally been associated with response inhibition (Aron et al., 2014), it also serves attentional control functions (Hampshire and Sharp, 2015) and theta band activity in this region has been connected to such processes (Wendiggensen et al., 2022). This is in line with other studies showing that theta band activity, as also found in sensory association areas likely serves attentional selection processes (Helfrich et al., 2018; VanRullen, 2018). It is, therefore, conceivable that the modulations of theta band activity in the observed brain regions by metacontrol biases reflect adjustments in attentional processes.

### Modulations of alpha and beta band dynamics

Alpha and beta frequency band activity was also affected by our metacontrol manipulation (Table 1, Figure 2). This suggests that metacontrol modulations are not confined to frequency bands assumed to play a key role in cognitive control, like theta, but can also impact frequency bands related to functions that are less directly associated with control. Interestingly, however, the modulations found for alpha and beta did not follow the pattern of the behavioral findings, as theta modulations did. This is possibly the case because the modulations of alpha and beta band activity occurred after the response was executed, which is particularly true for congruent trials. This already suggests that the role of alpha and beta band activity during metacontrol modulations might reflect the adjustment of control settings to optimize performance in the *next* trial. Both alpha and beta band activity were stronger during the FT_con_ than the FT_inc_ block, perhaps because control settings that are already rather optimal for dealing with response conflict, like the hypothesized persistence-biased setting in the FT_inc_ block, are harder to optimize even further. In contrast, rather relaxed, flexibility-biased settings like in the FT_con_, leave more operational space for increases of persistence. Note that this assumed scenario is consistent with predictions from conflict monitoring theory (Botvinick et al., 2001) and other control-adjustment approaches (Gratton et al., 1992; Keye et al., 2013).

Alpha band activity has been suggested to reflect inhibitory control processes regulating access of information of a knowledge system (Klimesch, 2011; Klimesch et al., 2007). Previous findings suggest that alpha band activity can re-instate after response selection processes have been finished and likely deactivate the previous stimulus-response association (Wolff et al., 2017). This is in line with the inhibitory function ascribed to alpha band activity (Klimesch, 2011; Klimesch et al., 2007). Corroborating our interpretation, the beamforming analysis (Table 1 and Figure 2) revealed that inferior frontal and superior frontal regions, as well as inferior/superior parietal regions were associated with alpha band activity modulations between the FT_inc_ and the FT_con_ blocks. Thus, regions known to exert a top-down control (i.e. frontal regions) (Aron et al., 2014; Miller and Cohen, 2001) are involved in the integrated processing of sensor and motor aspects during action selection (i.e. parietal regions) (Beste et al., 2023; Gottlieb, 2007; Gottlieb and Snyder, 2010; Wilken et al., 2023; Yu et al., 2024). The directed connectivity analysis revealed that for congruent trials (supplemental Figure S1 and Table 2), a bi-directional communication between all cortical regions was evident and the same was the case for the incongruent trials (supplemental Figure S2 and Table S3). Interestingly, the metacontrol manipulation by the experimental blocks (i.e. FT_con_ and FT_inc_) had an effect (Figures 3 and 4; Tables 2 and 3): the directed communication was always stronger in the persistence-heavy block (FT_inc_) than in the flexibility-heavy block (FT_con_). For congruent trials this referred to the connections from parietal to frontal regions (Figure 3) and for congruent trials this referred the connection between the superior frontal and ventral stream pathways (Figure 4).

The functional relevance of beta band activity is heavily debated (Barone and Rossiter, 2021; Engel and Fries, 2010; Kilavik et al., 2013; Spitzer and Haegens, 2017). Yet, some evidence suggest that post movement beta band activity (Pastötter et al., 2018) likely reflects the disintegration of stimulus-response associations used previously to guide action selection and responding, which lines up with evidence for content-specific BBA being able to change from active to latent to re-activated states (Spitzer and Haegens, 2017). The conducted beamforming analysis revealed that sensory, motor, and sensorimotor association cortices in frontal and parietal regions were associated with beta band activity modulations between the FT_con_ and the FT_inc_ block (Table 1, Figure 2). This corroborates the interpretation that the observed metacontrol-modulations of beta band activity may be related to a reset of stimulus-motor programs. The directed connectivity analysis revealed bi-direction communication between the involved cortical regions in congruent trials (supplemental Figure S1 and Table S1) and incongruent trials (supplemental Figure S2 and Table S2). This is line with an interpretation according to which system-level processes play a role. However, there was no significant effect between FT_inc_ and FT_con_ experimental blocks for the strength of directed connectivity (Figure 4 and Table 3). Directed communication in beta band associated cortical regions thus seems to be relevant for the reset regardless of whether a persistence-prone or a flexibility-prone bias was evident.

### Limitations and remarks

In almost every cognitive control task, including our Flanker task, decision-making processes are at play, involving both internally and externally guided components as distinguished by Nakao et al. (2016, 2012). Externally guided decision-making refers to situations where participants rely on external instructions to make a decision, such as selecting the correct key to press based on the presented stimuli—like in the Flanker task. Conversely, internally guided decision-making involves relying on internal strategies. For example, during a FT_con_ task block, participants’ performance is facilitated on congruent trials, suggesting they adopted an internal strategy to guide their decisions. From this perspective, we can assume that externally guided decisions imply or rely on a persistent metacontrol state that narrows attention to relevant external factors and maintains a strong commitment to externally defined rules or goals. Internally guided decisions, in turn, might promote a flexibility state that allows for broader, more exploratory responses to complex or ambiguous situations where the correct answer is not predefined. As outlined in the introduction, the functional role of theta band activity is likely closely connected to the alpha and beta band activity. It has been suggested that alpha band activity modulates theta band activity, which is turn may affect beta band activity during action control (Beste et al., 2023). Since the current findings revealed different neuroanatomical regions for the frequency bands, no interplay between these frequencies in terms of directed connectivity could be calculated. The current work intends to focus on oscillatory activity. Therefore, no aperiodic activity or fractal components were examined, known to be related to meta-control related aspects (Wainio-Theberge et al., 2022; Zhang et al., 2023). Future studies may also evaluate the impact of pre-stimulus activity, known to impact cognitive control processes (Wainio-Theberge et al., 2021; Wolff et al., 2021).

## Conclusion

In summary, we investigated how metacontrol states influence conflict regulation at a neurophysiological level by using EEG. We manipulated metacontrol states by varying the frequency of congruent and incongruent trials across experimental blocks and analyzed the effects on behavioral and electrophysiological measures. Behavioral data demonstrated a clear, predicted impact of the experimental manipulation. A persistence bias increased, and a flexibility bias reduced, conflict regulation. The EEG data, particularly in the theta frequency band, paralleled behavioral findings. Theta band activity modulations likely reflect the mismatch between expected metacontrol bias and actual task demands. Alpha and beta band dynamics also showed differences between experimental blocks, though these did not mirror behavioral effects. Alpha and beta activity, stronger when metacontrol was biased towards persistence, appeared post-response, indicating a system reset for upcoming demands. Directed connectivity analyses revealed stronger inter-regional communication, suggesting enhanced top-down control and sensorimotor integration in a persistence state. Theta band activity particularly reflects metacontrol processes, while alpha and beta bands are involved in resetting the system for future cognitive demands. These findings advance our understanding of the neural mechanisms underlying metacontrol and conflict monitoring.

## Acknowledgements

This work was supported by a Chinese Research Council (CSC) Fellowship awarded to X.W., and the Federal Ministry of Education and Research (Bundesministerium für Bildung und Forschung, BMBF) as part of the German Center for Child and Adolescent Health (DZKJ) under the funding code 01GL2405B.

## Notes

### Competing Interest Statement

The authors have declared no competing interest.

## References

Aron, A.R., Robbins, T.W., Poldrack, R.A., 2014. Inhibition and the right inferior frontal cortex: one decade on. Trends Cogn. Sci. (Regul. Ed.) 18, 177–185. 10.1016/j.tics.2013.12.003

Barone, J., Rossiter, H.E., 2021. Understanding the Role of Sensorimotor Beta Oscillations. Front. Syst. Neurosci. 15, 655886. 10.3389/fnsys.2021.655886

Beste, C., Baune, B.T., Falkenstein, M., Konrad, C., 2010. Variations in the TNF-α gene (TNF-α - 308G→A) affect attention and action selection mechanisms in a dissociated fashion. J. Neurophysiol. 104, 2523–2531. 10.1152/jn.00561.2010

Beste, C., Moll, C.K.E., Pötter-Nerger, M., Münchau, A., 2018. Striatal Microstructure and Its Relevance for Cognitive Control. Trends Cogn. Sci. (Regul. Ed.) 22, 747–751. 10.1016/j.tics.2018.06.007

Beste, C., Münchau, A., Frings, C., 2023. Towards a systematization of brain oscillatory activity in actions. Communications Biology 6. 10.1038/s42003-023-04531-9

Beste, C., Saft, C., Yordanova, J., Andrich, J., Gold, R., Falkenstein, M., Kolev, V., 2007. Functional compensation or pathology in cortico-subcortical interactions in preclinical Huntington’s disease? Neuropsychologia 45, 2922–2930. 10.1016/j.neuropsychologia.2007.06.004

Bigdely-Shamlo, N., Mullen, T., Kothe, C., Su, K.-M., Robbins, K.A., 2015. The PREP pipeline: standardized preprocessing for large-scale EEG analysis. Front. Neuroinform. 9. 10.3389/fninf.2015.00016

Botvinick, M.M., Braver, T.S., Barch, D.M., Carter, C.S., Cohen, J.D., 2001. Conflict monitoring and cognitive control. Psychol Rev 108, 624–652.

Buzsáki, G., 2006. Rhythms of the Brain. Oxford University Press.

Buzsáki, G., Draguhn, A., 2004. Neuronal oscillations in cortical networks. Science 304, 1926–1929. 10.1126/science.1099745

Cavanagh, J.F., Frank, M.J., 2014. Frontal theta as a mechanism for cognitive control. Trends Cogn. Sci. (Regul. Ed.) 18, 414–421. 10.1016/j.tics.2014.04.012

Chen, C.-C., Kilner, J.M., Friston, K.J., Kiebel, S.J., Jolly, R.K., Ward, N.S., 2010. Nonlinear Coupling in the Human Motor System. Journal of Neuroscience 30, 8393–8399. 10.1523/JNEUROSCI.1194-09.2010

Cifre, I., Miller Flores, M.T., Penalba, L., Ochab, J.K., Chialvo, D.R., 2021. Revisiting Nonlinear Functional Brain Co-activations: Directed, Dynamic, and Delayed. Front. Neurosci. 15, 700171. 10.3389/fnins.2021.700171

Cohen, M.X., 2014. A neural microcircuit for cognitive conflict detection and signaling. Trends Neurosci. 37, 480–490. 10.1016/j.tins.2014.06.004

Delorme, A., Makeig, S., 2004. EEGLAB: an open source toolbox for analysis of single-trial EEG dynamics including independent component analysis. Journal of Neuroscience Methods 134, 9–21. 10.1016/j.jneumeth.2003.10.009

Donoghue, T., Haller, M., Peterson, E.J., Varma, P., Sebastian, P., Gao, R., Noto, T., Lara, A.H., Wallis, J.D., Knight, R.T., Shestyuk, A., Voytek, B., 2020. Parameterizing neural power spectra into periodic and aperiodic components. Nat Neurosci 23, 1655–1665. 10.1038/s41593-020-00744-x

Elmers, J., Yu, S., Talebi, N., Prochnow, A., Beste, C., 2024. Neurophysiological effective network connectivity supports a threshold-dependent management of dynamic working memory gating. iScience 27, 109521. 10.1016/j.isci.2024.109521

Engel, A.K., Fries, P., 2010. Beta-band oscillations--signalling the status quo? Curr. Opin. Neurobiol. 20, 156–165. 10.1016/j.conb.2010.02.015

Ester, M., Kriegel, H.-P., Sander, J., Xu, X., 1996. A Density-Based Algorithm for Discovering Clusters in Large Spatial Databases with Noise 226–231.

Ferdousi, M., Babaie-Janvier, T., Robinson, P.A., 2020. Nonlinear wave-wave interactions in the brain. Journal of Theoretical Biology 500, 110308. 10.1016/j.jtbi.2020.110308

Friston, K.J., 2001. Book Review: Brain Function, Nonlinear Coupling, and Neuronal Transients. Neuroscientist 7, 406–418. 10.1177/107385840100700510

Gerster, M., Waterstraat, G., Litvak, V., Lehnertz, K., Schnitzler, A., Florin, E., Curio, G., Nikulin, V., 2022. Separating Neural Oscillations from Aperiodic 1/f Activity: Challenges and Recommendations. Neuroinform 20, 991–1012. 10.1007/s12021-022-09581-8

Goschke, T., Bolte, A., 2014. Emotional modulation of control dilemmas: the role of positive affect, reward, and dopamine in cognitive stability and flexibility. Neuropsychologia 62, 403–423. 10.1016/j.neuropsychologia.2014.07.015

Gottlieb, J., 2007. From thought to action: the parietal cortex as a bridge between perception, action, and cognition. Neuron 53, 9–16. 10.1016/j.neuron.2006.12.009

Gottlieb, J., Snyder, L.H., 2010. Spatial and non-spatial functions of the parietal cortex. Current Opinion in Neurobiology 20, 731–740. 10.1016/j.conb.2010.09.015

Gratton, G., Coles, M.G., Donchin, E., 1992. Optimizing the use of information: strategic control of activation of responses. J Exp Psychol Gen 121, 480–506. 10.1037//0096-3445.121.4.480

Hampshire, A., Sharp, D., 2015. Inferior PFC Subregions Have Broad Cognitive Roles. Trends Cogn. Sci. (Regul. Ed.) 19, 712–713. 10.1016/j.tics.2015.09.010

He, B.J., 2014. Scale-free brain activity: past, present, and future. Trends Cogn. Sci. (Regul. Ed.) 18, 480–487. 10.1016/j.tics.2014.04.003

He, F., Yang, Y., 2021. Nonlinear System Identification of Neural Systems from Neurophysiological Signals. Neuroscience 458, 213–228. 10.1016/j.neuroscience.2020.12.001

Helfrich, R.F., Fiebelkorn, I.C., Szczepanski, S.M., Lin, J.J., Parvizi, J., Knight, R.T., Kastner, S., 2018. Neural Mechanisms of Sustained Attention Are Rhythmic. Neuron 99, 854-865.e5. 10.1016/j.neuron.2018.07.032

Herrmann, C.S., Knight, R.T., 2001. Mechanisms of human attention: event-related potentials and oscillations. Neurosci Biobehav Rev 25, 465–476.

Hommel, B., 2015. Between Persistence and Flexibility, in: Advances in Motivation Science. Elsevier, pp. 33–67. 10.1016/bs.adms.2015.04.003

Hommel, B., 1994. Spontaneous decay of response-code activation. Psychol. Res 56, 261–268. 10.1007/BF00419656

Hommel, B., Colzato, L.S., 2017a. The social transmission of metacontrol policies: Mechanisms underlying the interpersonal transfer of persistence and flexibility. Neurosci Biobehav Rev 81, 43–58. 10.1016/j.neubiorev.2017.01.009

Hommel, B., Colzato, L.S., 2017b. Meditation and Metacontrol. J Cogn Enhanc 1, 115–121. 10.1007/s41465-017-0017-4

Hommel, B., Wiers, R.W., 2017. Towards a Unitary Approach to Human Action Control. Trends Cogn. Sci. (Regul. Ed.) 21, 940–949. 10.1016/j.tics.2017.09.009

Jia, S., Liu, D., Song, W., Beste, C., Colzato, L., Hommel, B., 2024. Tracing conflict-induced cognitive-control adjustments over time using aperiodic EEG activity. Cerebral Cortex 34, bhae185. 10.1093/cercor/bhae185

Keye, D., Wilhelm, O., Oberauer, K., Stürmer, B., 2013. Individual differences in response conflict adaptations. Front Psychol 4, 947. 10.3389/fpsyg.2013.00947

Kilavik, B.E., Zaepffel, M., Brovelli, A., MacKay, W.A., Riehle, A., 2013. The ups and downs of beta oscillations in sensorimotor cortex. Experimental Neurology 245, 15–26. 10.1016/j.expneurol.2012.09.014

Klimesch, W., 2012. Alpha-band oscillations, attention, and controlled access to stored information. Trends in Cognitive Sciences 16, 606–617. 10.1016/j.tics.2012.10.007

Klimesch, W., 2011. Evoked alpha and early access to the knowledge system: the P1 inhibition timing hypothesis. Brain Res. 1408, 52–71. 10.1016/j.brainres.2011.06.003

Klimesch, W., Sauseng, P., Hanslmayr, S., 2007. EEG alpha oscillations: the inhibition-timing hypothesis. Brain Res Rev 53, 63–88. 10.1016/j.brainresrev.2006.06.003

Kodama, N.X., Galán, R.F., 2019. Linear Stability of Spontaneously Active Local Cortical Circuits: Is There Criticality on Long Time Scales?, in: Tomen, N., Herrmann, J.M., Ernst, U. (Eds.), The Functional Role of Critical Dynamics in Neural Systems, Springer Series on Bio- and Neurosystems. Springer International Publishing, Cham, pp. 139–157. 10.1007/978-3-030-20965-0_8

Lai, M., Demuru, M., Hillebrand, A., Fraschini, M., 2018. A comparison between scalp- and source-reconstructed EEG networks. Sci Rep 8, 12269. 10.1038/s41598-018-30869-w

Logan, G.D., Zbrodoff, N.J., 1979. When it helps to be misled: Facilitative effects of increasing the frequency of conflicting stimuli in a Stroop-like task. Memory & Cognition 7, 166–174. 10.3758/BF03197535

Luck, S.J., Kappenman, E.S. (Eds.), 2013. The Oxford handbook of event-related potential components, Oxford library of psychology. Oxford University Press, Oxford ; New York, NY.

Mekern, V.N., Sjoerds, Z., Hommel, B., 2019. How metacontrol biases and adaptivity impact performance in cognitive search tasks. Cognition 182, 251–259. 10.1016/j.cognition.2018.10.001

Miller, E.K., Cohen, J.D., 2001. An integrative theory of prefrontal cortex function. Annu. Rev. Neurosci. 24, 167–202. 10.1146/annurev.neuro.24.1.167

Nakao, T., Kanayama, N., Katahira, K., Odani, M., Ito, Y., Hirata, Y., Nasuno, R., Ozaki, H., Hiramoto, R., Miyatani, M., Northoff, G., 2016. Post-response βγ power predicts the degree of choice-based learning in internally guided decision-making. Scientific Reports 6, 32477. 10.1038/srep32477

Nakao, T., Ohira, H., Northoff, G., 2012. Distinction between Externally vs. Internally Guided Decision-Making: Operational Differences, Meta-Analytical Comparisons and Their Theoretical Implications. Frontiers in Neuroscience 6. 10.3389/fnins.2012.00031

Nozari, E., Bertolero, M.A., Stiso, J., Caciagli, L., Cornblath, E.J., He, X., Mahadevan, A.S., Pappas, G.J., Bassett, D.S., 2020. Is the brain macroscopically linear? A system identification of resting state dynamics. 10.48550/ARXIV.2012.12351

Oostenveld, R., Fries, P., Maris, E., Schoffelen, J.-M., 2011. FieldTrip: Open source software for advanced analysis of MEG, EEG, and invasive electrophysiological data. Comput Intell Neurosci 2011, 156869. 10.1155/2011/156869

Papana, A., Kyrtsou, C., Kugiumtzis, D., Diks, C., 2013. Simulation Study of Direct Causality Measures in Multivariate Time Series. Entropy 15, 2635–2661. 10.3390/e15072635

Parra, L.C., Spence, C.D., Gerson, A.D., Sajda, P., 2005. Recipes for the linear analysis of EEG. NeuroImage 28, 326–341. 10.1016/j.neuroimage.2005.05.032

Pastötter, B., Engel, M., Frings, C., 2018. The Forward Effect of Testing: Behavioral Evidence for the Reset-of-Encoding Hypothesis Using Serial Position Analysis. Frontiers in Psychology 9. 10.3389/fpsyg.2018.01197

Pedroni, A., Bahreini, A., Langer, N., 2019. Automagic: Standardized preprocessing of big EEG data. NeuroImage 200, 460–473. 10.1016/j.neuroimage.2019.06.046

Pertermann, M., Bluschke, A., Roessner, V., Beste, C., 2019a. The Modulation of Neural Noise Underlies the Effectiveness of Methylphenidate Treatment in Attention-Deficit/Hyperactivity Disorder. Biol Psychiatry Cogn Neurosci Neuroimaging 4, 743–750. 10.1016/j.bpsc.2019.03.011

Pertermann, M., Mückschel, M., Adelhöfer, N., Ziemssen, T., Beste, C., 2019b. On the interrelation of 1/f neural noise and norepinephrine system activity during motor response inhibition. J. Neurophysiol. 121, 1633–1643. 10.1152/jn.00701.2018

Pi, Y., Yan, J., Pscherer, C., Gao, S., Mückschel, M., Colzato, L., Hommel, B., Beste, C., 2024. Interindividual aperiodic resting-state EEG activity predicts cognitive-control styles. Psychophysiology e14576. 10.1111/psyp.14576

Prochnow, A., Zhou, X., Ghorbani, F., Roessner, V., Hommel, B., Beste, C., 2024. Event segmentation in ADHD: neglect of social information and deviant theta activity point to a mechanism underlying ADHD. Gen Psych 37, e101486. 10.1136/gpsych-2023-101486

Roux, F., Uhlhaas, P.J., 2014. Working memory and neural oscillations: alpha–gamma versus theta– gamma codes for distinct WM information? Trends in Cognitive Sciences 18, 16–25. 10.1016/j.tics.2013.10.010

Ruiz-Gómez, S.J., Hornero, R., Poza, J., Maturana-Candelas, A., Pinto, N., Gómez, C., 2019. Computational modeling of the effects of EEG volume conduction on functional connectivity metrics. Application to Alzheimer’s disease continuum. J. Neural Eng. 16, 066019. 10.1088/1741-2552/ab4024

Schneider, T., Neumaier, A., 2001. Algorithm 808: ARfit—a matlab package for the estimation of parameters and eigenmodes of multivariate autoregressive models. ACM Trans. Math. Softw. 27, 58–65. 10.1145/382043.382316

Spitzer, B., Haegens, S., 2017. Beyond the Status Quo: A Role for Beta Oscillations in Endogenous Content (Re)Activation. eneuro 4, ENEURO.0170-17.2017. 10.1523/ENEURO.0170-17.2017

Talebi, N., Nasrabadi, A.M., Mohammad-Rezazadeh, I., Coben, R., 2019. nCREANN: Nonlinear Causal Relationship Estimation by Artificial Neural Network; Applied for Autism Connectivity Study. IEEE Transactions on Medical Imaging 38, 2883–2890. 10.1109/TMI.2019.2916233

Tzourio-Mazoyer, N., Landeau, B., Papathanassiou, D., Crivello, F., Etard, O., Delcroix, N., Mazoyer, B., Joliot, M., 2002. Automated Anatomical Labeling of Activations in SPM Using a Macroscopic Anatomical Parcellation of the MNI MRI Single-Subject Brain. NeuroImage 15, 273–289. 10.1006/nimg.2001.0978

Uhlhaas, P., 2009. Neural synchrony in cortical networks: history, concept and current status. Frontiers in Integrative Neuroscience 3. 10.3389/neuro.07.017.2009

Van Schependom, J., Baetens, K., Nagels, G., Olmi, S., Beste, C., 2024. Neurophysiological avenues to better conceptualizing adaptive cognition. Commun Biol 7, 626. 10.1038/s42003-024-06331-1

Van Veen, B.D., Van Drongelen, W., Yuchtman, M., Suzuki, A., 1997. Localization of brain electrical activity via linearly constrained minimum variance spatial filtering. IEEE Trans. Biomed. Eng. 44, 867–880. 10.1109/10.623056

VanRullen, R., 2018. Attention Cycles. Neuron 99, 632–634. 10.1016/j.neuron.2018.08.006

Voytek, B., Knight, R.T., 2015. Dynamic network communication as a unifying neural basis for cognition, development, aging, and disease. Biol. Psychiatry 77, 1089–1097. 10.1016/j.biopsych.2015.04.016

Wainio-Theberge, S., Wolff, A., Gomez-Pilar, J., Zhang, J., Northoff, G., 2022. Variability and task-responsiveness of electrophysiological dynamics: Scale-free stability and oscillatory flexibility. NeuroImage 256, 119245. 10.1016/j.neuroimage.2022.119245

Wainio-Theberge, S., Wolff, A., Northoff, G., 2021. Dynamic relationships between spontaneous and evoked electrophysiological activity. Commun Biol 4, 741. 10.1038/s42003-021-02240-9

Wendiggensen, P., Ghin, F., Koyun, A.H., Stock, A.-K., Beste, C., 2022. Pretrial Theta Band Activity Affects Context-dependent Modulation of Response Inhibition. Journal of Cognitive Neuroscience 34, 605–617. 10.1162/jocn_a_01816

Widmann, A., Schröger, E., Maess, B., 2015. Digital filter design for electrophysiological data – a practical approach. Journal of Neuroscience Methods 250, 34–46. 10.1016/j.jneumeth.2014.08.002

Wilken, S., Böttcher, A., Adelhöfer, N., Raab, M., Hoffmann, S., Beste, C., 2023. The neurophysiology of continuous action monitoring. iScience 26, 106939. 10.1016/j.isci.2023.106939

Willemssen, R., Müller, T., Schwarz, M., Falkenstein, M., Beste, C., 2009. Response monitoring in de novo patients with Parkinson’s disease. PLoS ONE 4, e4898. 10.1371/journal.pone.0004898

Winkler, I., Brandl, S., Horn, F., Waldburger, E., Allefeld, C., Tangermann, M., 2014. Robust artifactual independent component classification for BCI practitioners. J. Neural Eng. 11, 035013. 10.1088/1741-2560/11/3/035013

Winkler, I., Haufe, S., Tangermann, M., 2011. Automatic Classification of Artifactual ICA-Components for Artifact Removal in EEG Signals. Behav Brain Funct 7, 30. 10.1186/1744-9081-7-30

Wolff, A., Chen, L., Tumati, S., Golesorkhi, M., Gomez-Pilar, J., Hu, J., Jiang, S., Mao, Y., Longtin, A., Northoff, G., 2021. Prestimulus dynamics blend with the stimulus in neural variability quenching. NeuroImage 238, 118160. 10.1016/j.neuroimage.2021.118160

Wolff, N., Zink, N., Stock, A.-K., Beste, C., 2017. On the relevance of the alpha frequency oscillation’s small-world network architecture for cognitive flexibility. Sci Rep 7, 13910. 10.1038/s41598-017-14490-x

Yang, Y., Dewald, J.P.A., Van Der Helm, F.C.T., Schouten, A.C., 2018. Unveiling neural coupling within the sensorimotor system: directionality and nonlinearity. Eur J of Neuroscience 48, 2407–2415. 10.1111/ejn.13692

Yu, S., Konjusha, A., Ziemssen, T., Beste, C., 2024. Inhibitory control in WM gate-opening: Insights from alpha desynchronization and norepinephrine activity under atDCS stimulation. NeuroImage 289, 120541. 10.1016/j.neuroimage.2024.120541

Zhang, C., Beste, C., Prochazkova, L., Wang, K., Speer, S.P.H., Smidts, A., Boksem, M.A.S., Hommel, B., 2022. Resting-state BOLD signal variability is associated with individual differences in metacontrol. Sci Rep 12, 18425. 10.1038/s41598-022-21703-5

Zhang, C., Stock, A.-K., Mückschel, M., Hommel, B., Beste, C., 2023. Aperiodic neural activity reflects metacontrol. Cerebral Cortex 33, 7941–7951. 10.1093/cercor/bhad089

