## Supplemental for "Neurophysiological Dynamics of Metacontrol States: EEG Insights into Conflict Regulation"

### Congruent trials

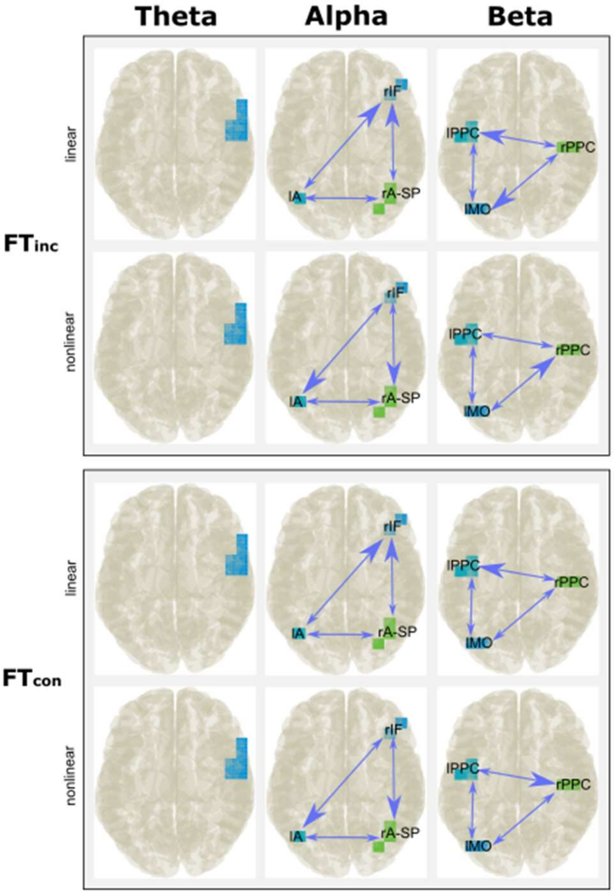


**Figure S1** The nCREANN Linear and Non-Linear connectivity between brain regions of congruent trials. The presented view is top view. Thicker arrow head indicate a higher connectivity after non-parametric tests.

**Table S1** Linear connectivity directions for congruent trial. The directions with significantly higher connectivity are marked in bold.

| $FT_{inc}$ | Theta | | | | |
|---|---|---|---|---|---|
| | Connectivity pairs | Median | Standard deviation | Z value | *p* value |
| | Single cluster | | | | |
| | Alpha | | | | |
| | **lA → rIF**<br>rIF → lA | Mdn = -1.5101<br>Mdn = -1.8329 | 0.2162<br>0.3203 | Z = -3.068 | *p* = .002 |
| | **rA-SP → rIF**<br>rIF → rA-SP | Mdn = -1.5043<br>Mdn = -1.7732 | 0.2557<br>0.2181 | Z = -3.346 | *p* < .001 |
| | lA → rA-SP<br>rA-SP → lA | | | | *ns* |
| | Beta | | | | |
| | lPPC → lMO<br>lMO → lPPC | | | | *ns* |
| | **rPPC → lMO**<br>lMO → rPPC | Mdn = -1.5843<br>Mdn = -1.7410 | 0.2970<br>0.2311 | Z = -2.705 | *p* = .007 |
| | **rPPC → lPPC** | Mdn = -1.4505 | 0.1785 | Z = -3.579 | *p* < .001 |

| | | | | | |
|---|---|---|---|---|---|
| | lPPC → rPPC | Mdn = -1.7453 | 0.2982 | | |
| FT$_{con}$ | Theta | | | | |
| | Single cluster | | | | |
| | Alpha | | | | |
| | **lA → rIF**<br>rIF → lA | Mdn = -1.6466<br>Mdn = -1.8990 | 0.2660<br>0.3230 | Z = -2.619 | *p* = .009 |
| | **rA-SP → rIF**<br>rIF → rA-SP | Mdn = -1.5888<br>Mdn = -1.7549 | 0.2351<br>0.2245 | Z = -2.057 | *p* = .040 |
| | lA → rA-SP<br>rA-SP → lA | | | | *ns* |
| | Beta | | | | |
| | lPPC → lMO<br>lMO → lPPC | | | | *ns* |
| | rPPC → lMO<br>lMO → rPPC | | | | *ns* |
| | **rPPC → lPPC**<br>lPPC → rPPC | Mdn = -1.5055<br>Mdn = -1.7580 | 0.2110<br>0.2688 | Z = -3.625 | *p* < .001 |

**Table S2** Nonlinear connectivity directions for congruent trial. The directions with significantly higher connectivity are marked in bold.

| FT$_{inc}$ | Theta | | | | |
|---|---|---|---|---|---|
| | Connectivity pairs | Median | Standard deviation | Z value | *p* value |
| | Single cluster | | | | |
| | Alpha | | | | |
| | **rIF → lA**<br>lA → rIF | Mdn = -1.7946<br>Mdn = -2.1072 | 0.5277<br>0.6001 | Z = -3.051 | *p* = .002 |
| | **rIF → rA-SP**<br>rA-SP → rIF | Mdn = -1.8495<br>Mdn = -2.2457 | 0.6924<br>0.4528 | Z = -2.300 | *p* = .021 |
| | lA → rA-SP<br>rA-SP → lA | | | | *ns* |
| | Beta | | | | |
| | lPPC → lMO<br>lMO → lPPC | | | | *ns* |
| | **lMO → rPPC**<br>rPPC → lMO | Mdn = -1.8177<br>Mdn = -2.3197 | 0.6857<br>0.6376 | Z = -2.064 | *p* = .039 |
| | rPPC → lPPC<br>lPPC → rPPC | | | | *ns* |
| FT$_{con}$ | Theta | | | | |
| | Single cluster | | | | |
| | Alpha | | | | |
| | **rIF → lA**<br>lA → rIF | Mdn = -1.7946<br>Mdn = -2.1072 | 0.5277<br>0.6001 | Z = -3.051 | *p* = .002 |
| | **rIF → rA-SP**<br>rA-SP → rIF | Mdn = -1.8495<br>Mdn = -2.2457 | 0.6924<br>0.4528 | Z = -2.300 | *p* = .021 |
| | lA → rA-SP<br>rA-SP → lA | | | | *ns* |
| | Beta | | | | |
| | lPPC → lMO<br>lMO → lPPC | | | | *ns* |
| | rPPC → lMO<br>lMO → rPPC | | | | *ns* |
| | **lPPC → rPPC**<br>rPPC → lPPC | Mdn = -1.7338<br>Mdn = -2.0542 | 0.4911<br>0.4753 | Z = -2.817 | *p* = .005 |

### Incongruent trials

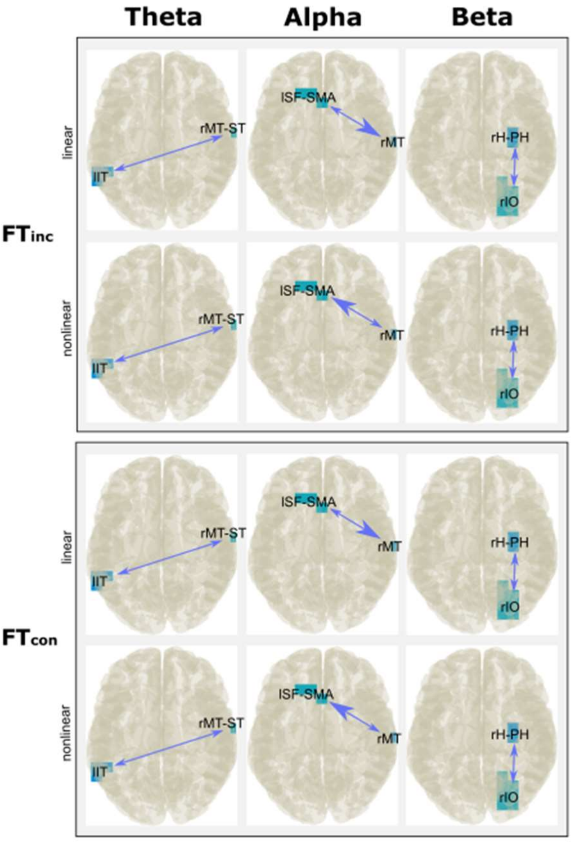


**Figure S2** The nCREANN Linear and Non-Linear connectivity between brain regions of incongruent trials. The presented view is top view. Thicker arrow head indicate a higher connectivity after non-parametric tests.

**Table S3** Linear connectivity directions for incongruent trial. The directions with significantly higher connectivity are marked in bold.

| $FT_{inc}$ | Theta | | | | |
|---|---|---|---|---|---|
| | Connectivity pairs | Median | Standard deviation | Z value | *p* value |
| | rMT-ST → lIT<br>lIT → rMT-ST | | | | *ns* |
| | Alpha | | | | |
| | **lSF-SMA → rMT**<br>rMT → lSF-SMA | Mdn = -1.6271<br>Mdn = -1.7585 | 0.1748<br>0.2473 | Z = -1.964 | *p* = .049 |
| | Beta | | | | |
| | rIO → rH-PH<br>rH-PH → rIO | | | | *ns* |
| $FT_{con}$ | Theta | | | | |
| | rMT-ST → lIT<br>lIT → rMT-ST | | | | *ns* |
| | Alpha | | | | |
| | **lSF-SMA → rMT**<br>rMT → lSF-SMA | Mdn = -1.4983<br>Mdn = -1.7906 | 0.1479<br>0.1979 | Z = -3.680 | *p* < .001 |
| | Beta | | | | |
| | rIO → rH-PH<br>rH-PH → rIO | | | | *ns* |

**Table S4** Nonlinear connectivity directions for incongruent trial. The directions with significantly higher connectivity are marked in bold.

| $FT_{inc}$ | Theta | | | | |
|---|---|---|---|---|---|
| | Connectivity pairs | Median | Standard deviation | Z value | *p* value |

| | | | | | |
|---|---|---|---|---|---|
| | rMT-ST → lIT<br>lIT → rMT-ST | | | | *ns* |
| | Alpha | | | | |
| | **rMT → lSF-SMA**<br>lSF-SMA → rMT | Mdn = -1.7962<br>Mdn = -2.2749 | 0.4379<br>0.6095 | Z = -3.411 | $p < .001$ |
| | Beta | | | | |
| | rIO → rH-PH<br>rH-PH → rIO | | | | *ns* |
| $FT_{con}$ | Theta | | | | |
| | rMT-ST → lIT<br>lIT → rMT-ST | | | | *ns* |
| | Alpha | | | | |
| | **rMT → lSF-SMA**<br>lSF-SMA → rMT | Mdn = -1.6647<br>Mdn = -2.1840 | 0.3475<br>0.4795 | Z = -4.056 | $p < .001$ |
| | Beta | | | | |
| | rIO → rH-PH<br>rH-PH → rIO | | | | *ns* |
